# FIDDLE: An integrative deep learning framework for functional genomic data inference

**DOI:** 10.1101/081380

**Authors:** Umet Eser, L. Stirling Churchman

## Abstract

Numerous advances in sequencing technologies have revolutionized genomics through generating many types of genomic functional data. Statistical tools have been developed to analyze individual data types, but there lack strategies to integrate disparate datasets under a unified framework. Moreover, most analysis techniques heavily rely on feature selection and data preprocessing which increase the difficulty of addressing biological questions through the integration of multiple datasets. Here, we introduce FIDDLE (Flexible Integration of Data with Deep LEarning) an open source data-agnostic flexible integrative framework that learns a unified representation from multiple data types to infer another data type. As a case study, we use multiple *Saccharomyces cerevisiae* genomic datasets to predict global transcription start sites (TSS) through the simulation of TSS-seq data. We demonstrate that a type of data can be inferred from other sources of data types without manually specifying the relevant features and preprocessing. We show that models built from multiple genome-wide datasets perform profoundly better than models built from individual datasets. Thus FIDDLE learns the complex synergistic relationship within individual datasets and, importantly, across datasets.

## Introduction

Functional annotation of a particular genomic region requires probing multiple molecular interactions under different conditions. The combinatorial complexity of genomic interactions and various perturbative conditions dictate that probing each one-by-one is extremely laborious and resource prohibitive. Thus, there is a critical need for integrative approaches that learn the unified representation of a given genomic region from heterogeneous datasets to infer data that has not been directly obtained. Furthermore, high-level questions are best answered by the evaluation of multiple genomic datasets simultaneously and will benefit from a unifying easy-to-use integrative framework.

Genomics data provide position specific information about molecular interactions that take place on the DNA. For a number of cell types numerous datasets are publicly available in large databases, such as ENCODE and Roadmap Epigenome (The ENCODE Project Consortium 2012; Roadmap Epigenomics Consortium et al. 2015). It is highly challenging to integrate these datasets to learn a unified representation for a given task. First of all, these datasets are often highly dimensional. In other words, a genomic region of interest is represented many numbers across the region that arise from experiments that make measurements for each basepair. Second, the data to be integrated have different spatial resolution and can be discrete or continuous valued. Last but not least, such data comprise complex structures, *i.e.* a position can correlate with other positions within or across datasets. Current integrative approaches are highly customized as they depend on domain knowledge, data pre-processing and feature selection, which creates a bottleneck for learning rich representations in a reusable and transferable manner.

Recently, in the fields of computer vision (Krizhevsky, Sutskever, and Hinton 2012) and natural language processing (Collobert et al. 2011), a breakthrough has been achieved by automatic learning of features solely from data using deep learning. Convolutional neural networks (ConvNets) were utilized to adaptively learn the features instead of choosing manually.ConvNets nonlinearly transforms the data to decouple the complex internal structure that arises from the correlation between the low level features and provide an informative rich representations that simplifies classification or regression (Bengio 2013). Several studies have adopted ConvNets to DNA sequence analysis to predict protein binding, protein contact map, alternative splicing and accessibility from DNA sequence (Alipanahi et al. 2015; Xiong et al.2015; Kelley, Snoek, and Rinn 2016; Wang et al. 2016). Moreover, as in many other fields, deep learning approaches are shown to outperform classical well-known machine learning algorithms such as support vector machines and random forests (Rusk 2015; Angermueller et al. 2016).Deep learning has the potential to statistically learn unified rich representations from multiple data sources (modalities) (Castrejon, Aytar, and Vondrick, n.d.), yet has not been readily accessible by experimental biologists due to technical and notational reasons.

Here, we provide an open source package, called FIDDLE (https://github.com/ueser/FIDDLE), that comprises ConvNet modules for individual datasets and combines under a common scaffold for unified data representation for dataset inference. We used FIDDLE to infer yeast Transcription Start Site sequencing (TSS-seq) (Malabat et al. 2015) data from NET-seq (Churchman and Weissman 2011), MNase-seq, TFIIB-ChIP-seq, RNA-seq (Hughes et al. 2012) and DNA sequence, individually and combined. We first demonstrate that these individual data types contain information about TSS-seq by quantifying the prediction of conditional probability distribution and peak detection accuracy. We then evaluate the performance by comparing the prediction from biological replicates as upper limit. Next, we show that the combined model that uses all of the datasets synergistically boosts the performance towards the upper limit of accuracy. Finally, we dissect the contribution of individual datasets by defining necessity and sufficiency scores. We find that DNA sequence alone is not sufficient and produces the model with the least accurate predictions, however, when combined with the other data, it becomes necessary to maintain the high accuracy. Such a synergistic relationship denotes the importance of data integration through automatic feature learning. As a result, FIDDLE provides a flexible, data agnostic framework to integrate multiple data types and infer a dataset.

## Results

Genomics data originate from experiments designed to target specific molecular interactions, however, they are likely to indirectly provide information about other biological processes. For instance, Native Elongating Transcript Sequencing (NET-seq) directly monitors the distribution of actively transcribing polymerase along the genome, but also indirectly informs about the transcription initiation, nucleosome occupancy, transcript levels, splicing activity etc. (Churchman and Weissman 2011). However, it is highly challenging to exploit such secondary information as it requires knowing the relevant features beforehand. One of the premises of deep learning is that it enables automatic extraction of relevant features at multiple scales.

Therefore, we built a generic ConvNet module with 2 convolutional layers and a fully connected layer using Tensorflow (Figure 1a). Each layer applies a non-linear affine transformation to the tensor produced by the previous layer and conducts the transformed tensor to the next layer.Output of this ConvNet module is a representation of the input data which is concatenated with the representations of other datasets. Then, an additional convolutional layer is applied to the concatenated representations to learn the patterns emerge from synergistic interactions between input datasets. Finally, a fully connected layer is applied to the output of the combined convolution layer to predict the non-parametric probability distribution of the target. (Figure 1b).

**Figure 1.**
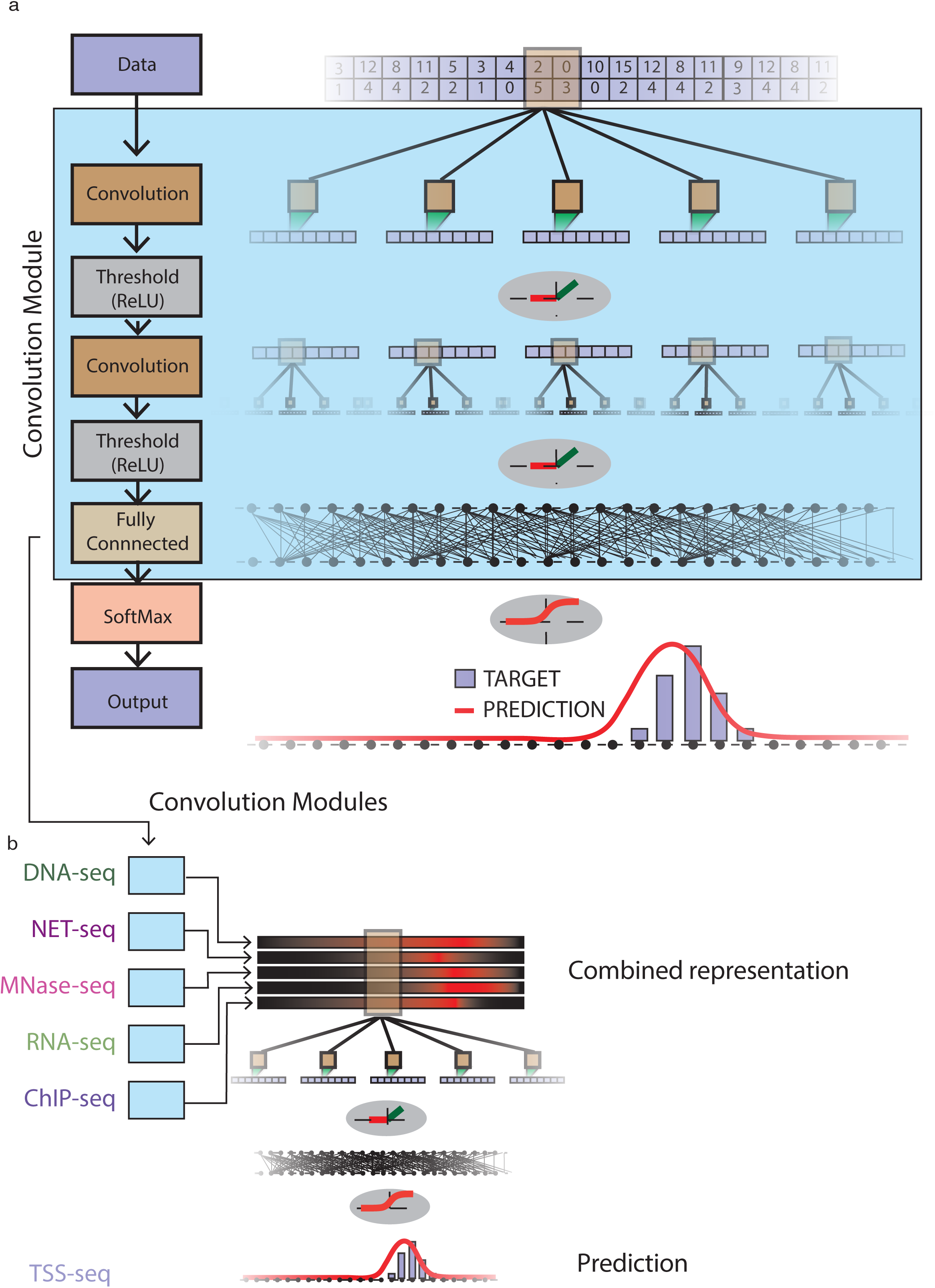
Schematic of FIDDLE. **(a)** FIDDLE integrates convolution modules that are dedicated
to learn the specific features of individual input types. A generic convolution module has two
convolutional layers each of which learns the patterns through the convolutional filters in different length scales. **(b)** An additional convolutional layer over the combined representation learns the patterns that appear across data types.

The raw sequencing data are mapped to DNA on both strands, even if the experiment does not have any strand specificity, such as MNase-seq and ChIP-seq. Usually during alignment both strands are aggregated into one track and are shifted by a certain amount or by a sophisticated analysis such as template filtering (Weiner et al. 2010). FIDDLE neither requires aligning the signal by shifting, nor needs any peak detection to find transcription factor or nucleosome positioning. It takes both strands as two tracks of an input to a ConvNet module. Moreover, to have a readily usable universal framework, we avoided input dependent preprocessing such as denoising, normalizing etc. and directly used the mapped sequencing data as inputs. When using DNA sequence as an input, we do not provide any pre-defined DNA sequence motifs. We convert DNA sequence into a binary matrix of size 4xL by one-hot-encoding, where L is the length of the DNA sequence.

Throughout the study, we prepared the inputs and output (TSS-seq) as L=500bp regions taken from a larger region that spans from 1 kp upstream and 1 kp downstream of non-overlapping genes’ start sites, producing 129K samples (see example in Figure 2).Then we split our data into 128K training and 1K testing. As our aim is to provide a universally flexible and readily available module, we chose reasonable hyperparameters as suggested previously and did not optimize them for this particular task (Angermueller et al. 2016). However, to see whether FIDDLE is sensitive to hyperparameters, we performed experiments by changing batch size, learning rate, number and size of convolution filters. We observe only slight changes on performance.

**Figure 2.**
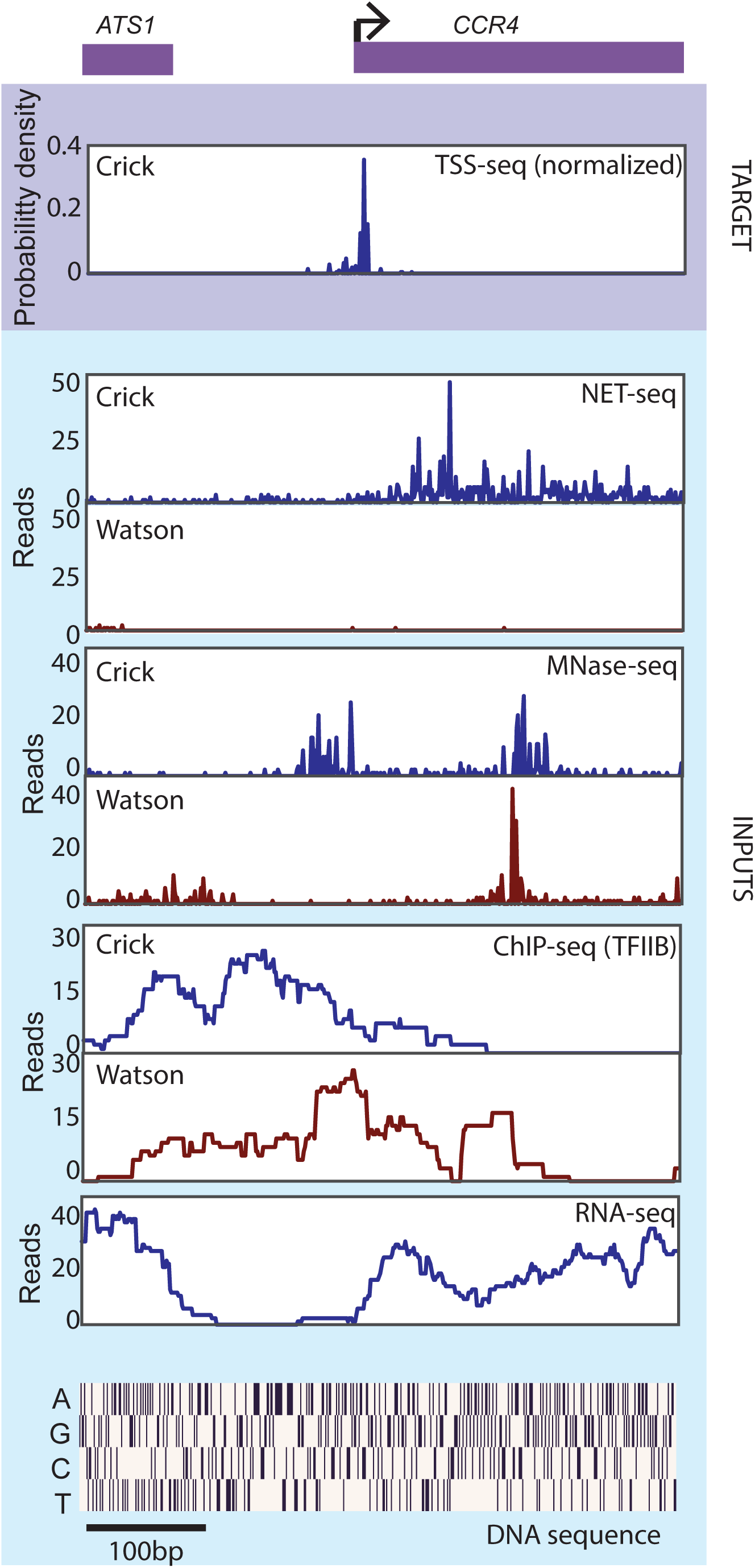
Example of training data at the *CCR4* locus. Probability density estimate of the target (TSS-seq), raw counts of the inputs, NET-seq, MNase-seq, TFIIB ChIP-seq and RNA-seq, and the one-hot-encoded representation of DNA sequence corresponding to the 500bp region that entails the up- and down-stream of *CCR4* gene promoter are shown. Reads from NET-seq, MNase-seq and ChIP-seq are strand specific and labeled (Watson or Crick).

To demonstrate that a dataset can be inferred from other data types, we trained our ConvNet modules to predict TSS-seq data for individual input datasets. All of them converge immediately to a plateau practically within the first epoch, in other words, before the whole dataset has been seen by the model (Figure 3b). It is worth noting that when DNA sequence is used as the input, it takes slightly more iteration to converge to a plateau. This is probably due to the digital nature of the DNA sequence, which informs the probability distribution of TSS via motifs. Although learning from NET-seq is slightly slower than MNase-seq, it asymptotically outperforms the other inputs possibly because NET-seq provides single nucleotide resolution signal whereas the resolution of other datasets are broader. Together, these results suggest that a dataset can be derived from other datasets, even without specifying the high level features.

**Figure 3.**
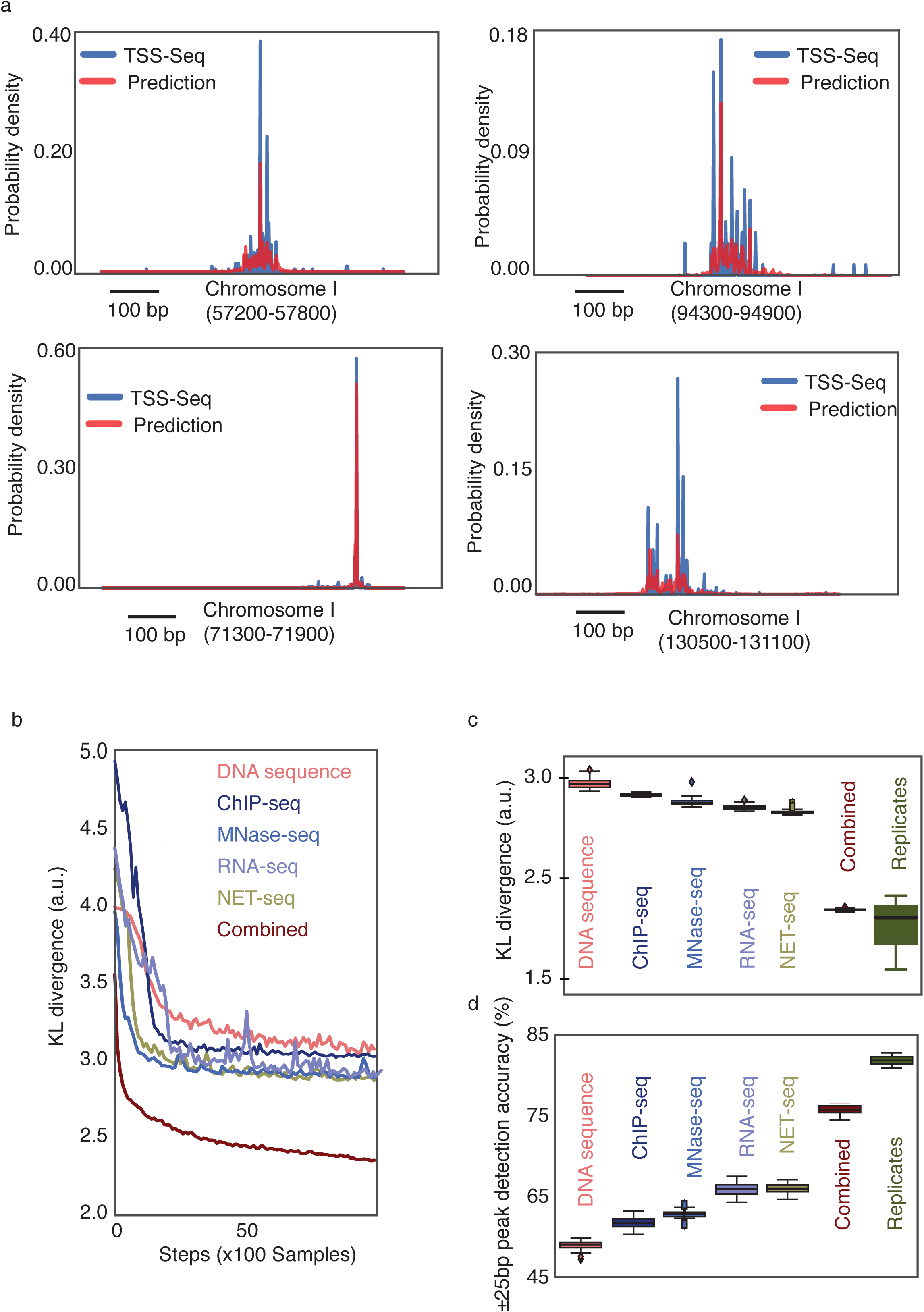
Evaluation of FIDDLE performance. **(a)** Examples of TSS-seq data and model predictions. FIDDLE learns arbitrary non-parametric probability distributions. Besides capturing overall shape of the distribution, the predictions can also emphasizes the major TSS site at nucleotide resolution. **(b)** Learning curve denotes the decrease of the Kullback-Leibler divergence between the prediction and the target by the number of held-out samples. **(c)** Summary statistics of the KL-divergence and the accuracy of the held-out data for individual data types, combined training and the ones measured from biological replicates.

Output of the model is a probability distribution over 500 bp genomic region, denoting the transcription initiation probability at each nucleotide, given that transcription initiates within the region. To evaluate how well the individual ConvNet modules perform, we compared the Kullback-Leibler divergence (KL divergence) between the prediction and the TSS-seq data, both of which approximate the probability distribution of the presence of a TSS over the 500 bp region, given there is at least one TSS in the same region (Figure 3c). As it is difficult to interpret KL-divergence, we also devised a metric to intuitively compare the accuracy of the predictions. Although the TSS is distributed over a region, in many bioinformatics applications, major transcription start site (peak position of the distribution) is sufficient. Therefore, we first divided 500 bp region into 10 non-overlapping bins, where each bin is a 50bp window. If both the model prediction and the TSS-seq data have their maxima within the same bin, we called the prediction correct. Randomized input-output datasets serve as a lower limit (10.5%) and biological replicates of TSS-seq data provide an estimate of the upper limit (81.2%) for accuracy for this type of measurements. We observe that NET-seq achieves 61.2% and the rest achieve the following accuracies: RNA-seq 60.1%, MNase-seq 55.4%, TFIIB ChIP-seq 53.9% and DNA sequence 50.2%. (Figure 3d)

To see whether the prediction performance is further improved when the model is trained with all input datasets, we combined the ConvNet modules under the scaffold module. Then we trained the combined model by randomly initializing the parameters. As expected, the combined model achieves significantly higher performance than the individually trained models (72.6%,Figure 3c,d). The combined FIDDLE model is able to predict arbitrary non-parametric probability distributions (Figure 3a). Note that the model prediction can adopt the major transcription start site preference at the nucleotide scale from the combination of inputs many of which have lower resolutions. These results suggest that the information to predict TSS-seq is not redundant across different datasets.

As the combined model radically improves performance, we hypothesized that the model learns the synergistic relationship between datasets when predicting the TSS. In other words, the model may utilize the information within a specific input dataset differently depending on the other input datasets’ context. For example, the combined model may ignore the presence of a DNA sequence motif within a specific region, if the corresponding NET-seq signal indicates no transcriptional activity. One way to understand whether there is a synergistic structure across datasets, is to measure the individual contribution of the input datasets to the combined model prediction. If a dataset is not sufficient but necessary to achieve high accuracy, then we deduce that the model is able to learn synergistic behavior. Therefore, we defined two scores, namely necessity and sufficiency. Necessity is determined by setting the input to its mean value over samples and measuring the cost discrepancy from the unperturbed prediction. If cost increases significantly, the necessity score will be high, which suggests that the input is necessary for accurate prediction. On the other hand, if the particular input is redundant given other inputs, we expect that the cost will not change significantly and therefore will have low necessity score. Similarly, the sufficiency score is calculated by setting all inputs but the one of interest to their sample mean. Then we compare the cost with the unperturbed model. If the isolated input is alone sufficient to accurately predict the output, we do not expect to see a significant change in the cost and vica versa. To control for the sufficiency score, in another model, we also provide TSS-seq as input as it is evident that a dataset is sufficient to predict itself. Likewise, to control for the necessity score, we provide white noise as input which does not contain any information about TSS-seq and is not necessary at all. Overall, many of the datasets have high sufficiency scores and low necessity scores, implying that they each contain strong secondary information about where the TSS are (Figure 4). An interesting exception is the DNA sequence data.Interestingly, we observe that the DNA sequence is necessary to maintain the high accuracy but not sufficient by itself. This suggests that the model utilizes the information from DNA sequence in combination with the other data types to accurately infer the TSS probability distribution.

**Figure 4.**
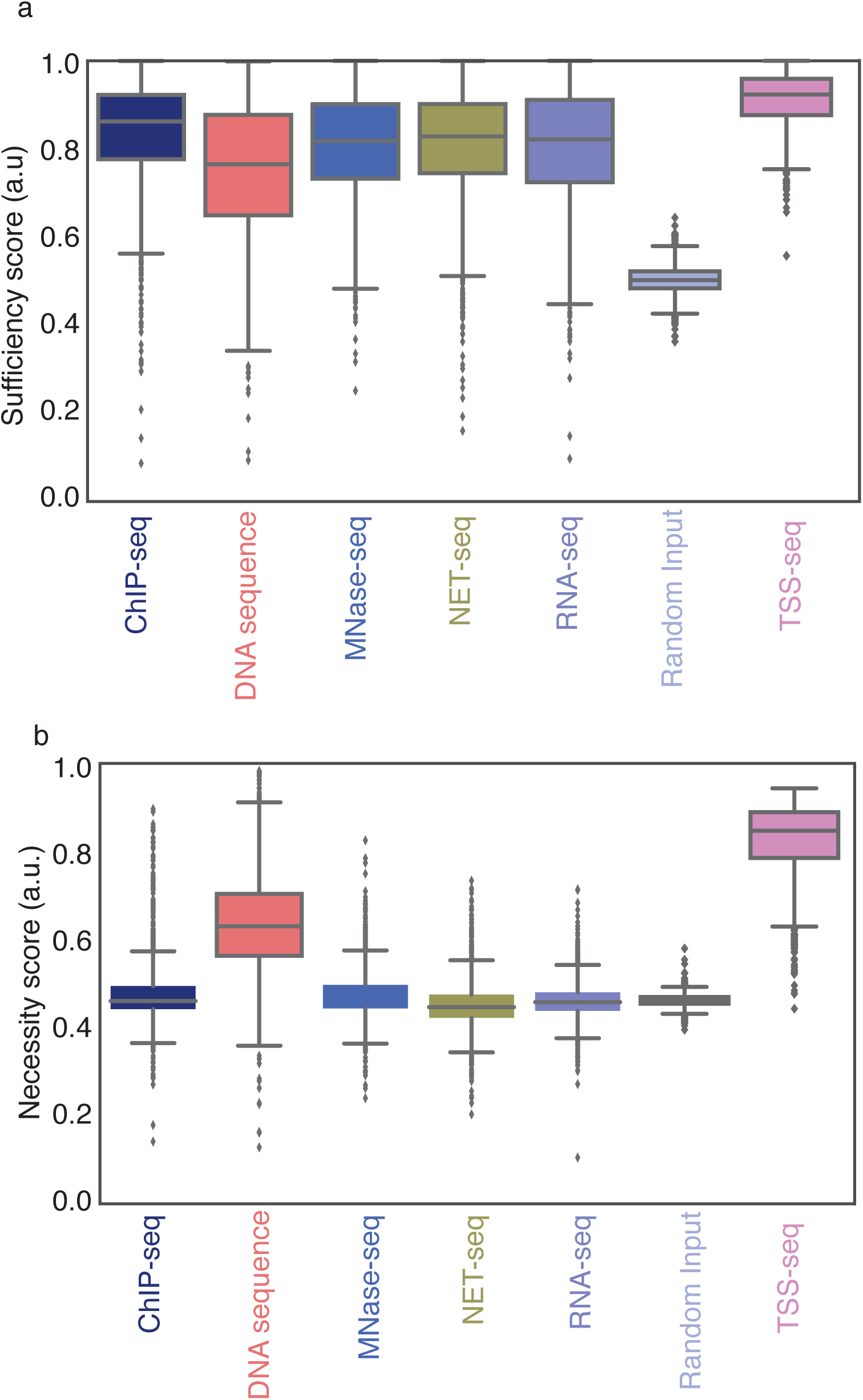
FIDDLE uses conditional information across input data types. Summary statistics of sufficiency score **(a)** and necessity score **(b)** for randomly selected 2000 data points.

## Discussion

In this study, we introduce FIDDLE as a Flexible Integration of Data with Deep LEarning for data inference. We demonstrate that the TSS-seq data can be learned accurately from different data types individually, and from their combination. The performance was boosted when the datasets are combined, because the FIDDLE can learn the context dependent synergistic relationship between datasets.

Although the performance of FIDDLE is already high, further improvement could be obtained by optimizing the hyperparameters, such as number of layers. Moreover, the crafting the ConvNet module specifically to the input type would likely improve performance. However, we have not designed specific custom modules so that FIDDLE is as data agnostic as possible to encourage general use.

We also avoid hyperparameter search and data preprocessing. The dynamic range of certain datasets (such as NET-seq) spans several order of magnitudes, which is usually worrisome for convergence and model generalization. However, with batch normalization (Ioffe and Szegedy 2015), the internal covariate shift is reduced and a large dynamic range becomes less effective.

FIDDLE is a generic framework that learns a unified rich representation by exploiting synergistic interactions within and across datasets. This representation can then be used to infer a particular dataset. Clearly with the cost and widespread need of genome-wide datasets, the potential of this framework is vast. All that is required for predicting a dataset is to constrain the model through aligning the representations of inputs within a context specified by the data to be predicted. Different representations can be learned by specifying different contexts. For example, we could ask FIDDLE to predict transcription factor binding sites using the same input datasets. In this case, the model would learn a different unified representation through the new constraints. These representations can then be used in a number of flexible ways, such as to infer causality between genomic features or to transfer the relationship to another domain, such as another species or cell type where the target dataset is not available.

## Methods

### Model architecture and parameters

Each convolutional layer also contains an average pooling, rectified linear unit, which thresholds negative values, and a batch normalizer unit. Batch normalization is shown to be important for fast convergence and better performance (Ioffe and Szegedy 2015). This generic convolutional module accepts input as H by L matrices and outputs 1x500 vector, where H is the number genomic data tracks and L is the length of the region of interest for a specific dataset. For the first convolution layer, we used 80 filters with size Hx5, where H is the height of the input matrix, depends on the input (e.g., NET-seq has 2x5, DNA sequence has 4x5 etc.). For the second convolution layer, we used 40 of 1x5 convolution filters. Finally, there is a scaffold module which concatenates the output vectors of individual ConvNet modules into a matrix of size Nx500, where N is the number of inputs.

The scaffold module contains 1 convolution layer which has an average pooling, ReLU and batch normalizer unit. Specifically, we used 20 of Nx5 convolution filters for the scaffold convolution layer. The final layer is the unified representation of all datasets to predict TSS-seq data. We then apply logistic regression to predict the TSS-seq signal of size 1xL for the same region of interest as a probability distribution. The target data, TSS-seq, is converted to probability distribution by dividing the signal to total reads for the particular region. Therefore, the overall model predicts the conditional probability distribution of transcription start site, conditioned on the region. We use Kullback-Leibler divergence as a cost function to train FIDDLE.

### Description of datasets

Native Elongating Transcript Sequencing (NET-seq) monitors actively elongating RNA Polymerase II by sequencing the 3’ ends of the associated nascent RNA. It provides nucleotide resolution data as the position of the 3’ end of the aligned reads denotes the RNAPII active site position. NET-seq also captures the unstable transcripts which might provide extra information about the queried system. The dataset is obtained from Churchman and Weissman, 2011. For every sample point, we concatenate NET-seq vectors of the positive and negative strands of size 500 into a 2x500 matrix and provide as the input.

Micrococcal Nuclease Sequencing (MNase-seq) captures the nucleosome occupancy by sequencing the DNA that is protected from nuclease digestion by nucleosomes (Hughes et al. 2012). MNase-seq dataset is not strand specific and usually the signals on the strands are shifted and merged. However, instead of merging the signal after processing, we make 2x500 matrices as input samples and leave the feature specification to FIDDLE.

TFIIB-Chromatin ImmunoPrecipitation Sequencing (ChIP-seq) probes the TFIIB protein along the DNA by standard ChIP-seq procedure (Hughes et al. 2012). The dataset is not strand specific, and have a resolution around ~100bp. As we do for other datasets, we provide the reads from both strands without aligning to each other as an input, resulting a 2x500 matrix for a data point.

RNA Sequencing (RNA-seq) measures the amount of transcripts by sequencing RNA. RNA-seq can be strand specific but in this case, we use unspecific one (Hughes et al. 2012). In this case, we merge both strands as it does not require strand alignments and use the spatial genomic profile.

Transcription Start Site Sequencing (TSS-seq) is a modified version of 5’ RACE, which captures 5’ end capped transcripts and sequences (Malabat et al. 2015). When mapped to the genome, only 5’ end is retained, denoting the TSS with a nucleotide resolution. We use this dataset as the target to be predicted. Instead of predicting unbounded reads, we converted the signal into probability density by dividing the reads over region of interest by the total read within the same region.

### Accuracy calculation

To calculate the accuracy, we divided the 500 bp region into 10 non-overlapping bins. Then we counted the number of times that peaks of the model prediction and the TSS-seq data fall into the same bin. This is a conservative score as the peaks that fall into neighboring bins are considered as incorrect regardless of their proximity. However, there may be local maximum of the TSS signal in the predicted bin. Similarly, we calculated the accuracy of biological replicates by counting the percentage of the data points that the peaks of the both replicates fall into the same bin.

### Sufficiency and necessity scores

To understand whether the model utilizes the interdependencies across input datasets, we defined two scores, namely sufficiency and necessity.

Let *L*(*x*) be the loss of the combined model,*L*_*d*(*x*) be the loss of the combined model when the dataset *d* is set to its sample mean and *L*+*d*(*x*) be the loss of the combined model when all other datasets are set to their sample mean but *d*, for a datapoint *x*. Then the sufficiency score *S* and necessity score *N* are defined by,

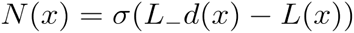

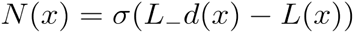

where *σ* is the sigmoid function, applied to bound the scores between 0 and 1.

## Acknowledgements

We would like to thank Yusuf Aytar and Ilker Yildirim for their useful discussions on robust training of deep learning models for multimodal data. This work was supported by US National Institutes of Health NHGRI grants R01HG007173 to L.S.C. and a Burroughs Wellcome Fund Career Award at the Scientific Interface to L.S.C.

